# Chromosomal requirements for formation of multivalent human nucleoli

**DOI:** 10.64898/2026.06.08.730829

**Authors:** Krystyna Giemza, Hazel Mangan, Brian McStay

**Author notes:** co-first authors (Krystyna Giemza, Hazel Mangan).

## Abstract

Human nucleoli are multivalent, involving contributions from up to ten NOR-bearing acrocentric chromosome p-arms. Precision mega-base scale chromosome engineering defines the requirements for this major genome organisational event. NOR deletions reveal that p-arm nucleolar association is rDNA independent. Deletion of all NOR-distal or proximal sequences individually have only a marginal effect on nucleolar association. Finally, deletion of an entire p-arm, while leaving centromere function intact, destroys the nucleolar association potential of that acrocentric. We propose that formation of multivalent nucleoli is not a nucleolar fusion event *per se;* rather it is driven by the surrounding chromosomal context of NORs.

## Introduction

Human chromosomes occupy distinct non-randomly positioned territories and are folded into hierarchical domains to facilitate genome packaging and organisation of functional compartments (Misteli 2020). Nucleoli, the most prominent functional compartments, form around arrays of ribosomal genes (rDNA) known as nucleolar organiser regions (NORs) located on the short p-arms of the five acrocentric chromosomes 13, 14, 15, 21 and 22 (Henderson et al. 1972; McStay 2023).

In non-transformed human cells the default status of NORs is active as defined by the presence of UBF, a nucleolar-specific HMG-box DNA binding protein that binds extensively across the rDNA array (van Sluis et al. 2020). Cells could in principle contain 10 nucleoli, yet most have between one and three, each involving multiple acrocentric p-arms, organised into discrete NOR territories with associated peri-nucleolar heterochromatin (PNH) (Mangan and McStay 2021; Leeke et al. 2025). We propose referring to these as ‘multivalent nucleoli’, in the sense that multiple acrocentric p-arms are involved in their formation, but no mechanism implied. Thus, formation of multivalent nucleoli involves appropriate juxtaposition of multiple chromosome territories and hierarchical domains within them, representing a major genome organisation event.

Human cells typically contain between 300 and 600 rDNA repeats, unevenly distributed among the five pairs of acrocentric p-arms (Stults et al. 2008; van Sluis et al. 2020; Potapova et al. 2025). NORs are bordered on their telomeric and centromeric sides by the distal junction (DJ) and proximal junction (PJ) respectively, shared among the acrocentrics (Fig. 1A) (Floutsakou et al. 2013; van Sluis et al. 2019). The DJ (∼ 300kb) is dominated by a large (>100kb) inverted repeat, which encodes long non-coding RNAs (lncRNAs) required for NOR function. 3D-immuno FISH with a DJ FISH probe has proved a reliable method for enumerating p-arms in each nucleolus. The arrival of Telomere to Telomere (T2T) genomes has delivered full sequences for acrocentric p-arms, heralding a new era of research into how they function (Nurk et al. 2022; Guarracino et al. 2023). They range in size from 10-17Mb with NOR distal and proximal sequences comprising multiple satellite DNA classes and segmental duplications (SDs).

**Figure 1.**
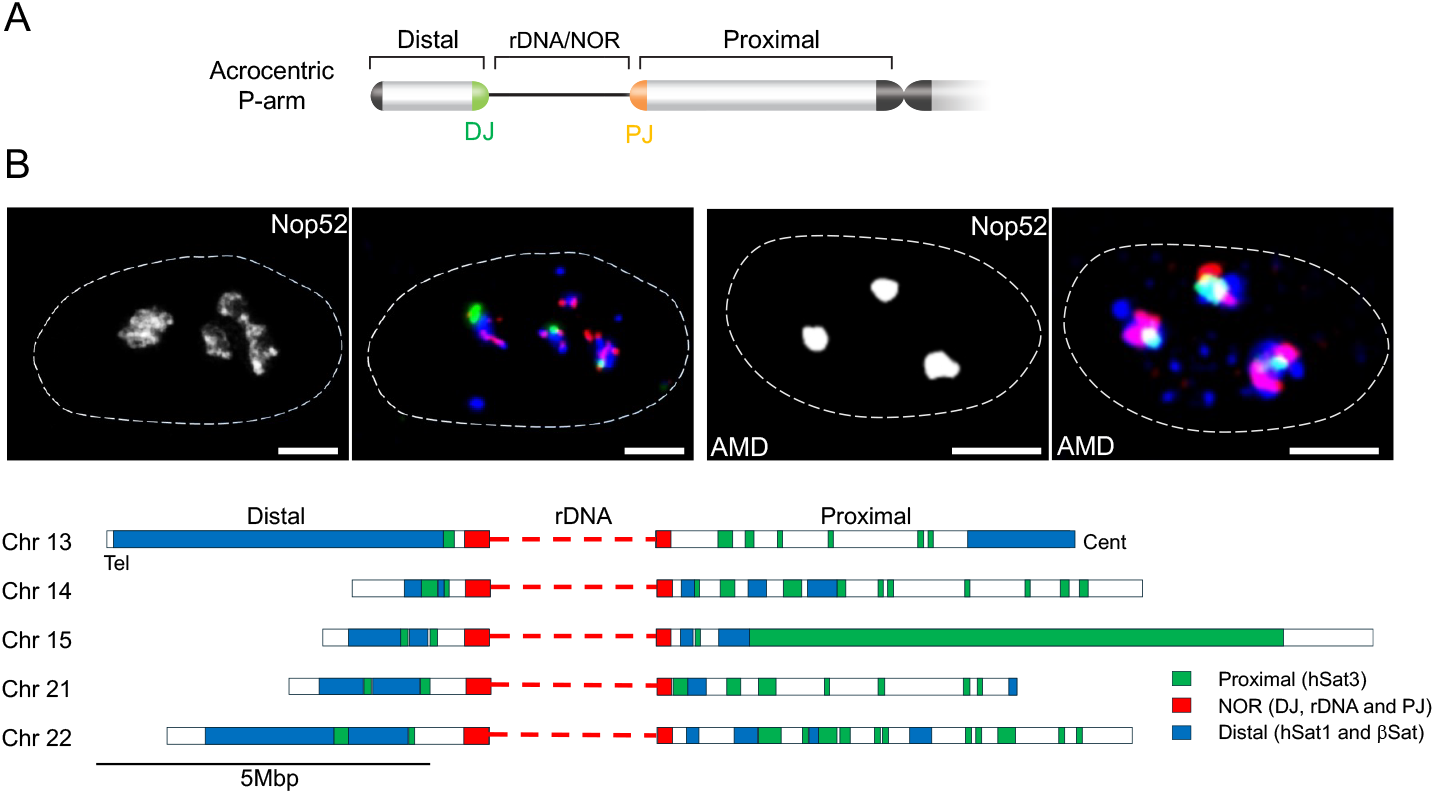
Acrocentric p-arms associate with nucleoli. (A) Organization of NORs on acrocentric p-arms, depicting the NOR, distal and proximal regions, distal and proximal Junctions (DJ and PJ) are depicted in green and orange respectively. (B) 3D-immunoFISH was performed on untreated (left panels) and AMD treated (right panels) RPE1 cells with distal, NOR and proximal p-arm regional probes and antibodies against Nop52. Nuclear membranes are indicated by dashed white lines and scale bars represent 5mm. Probe compositions and the distribution of their target sequences on acrocentric p-arms in T2T CHM13 are shown below.

In the past, formation of multivalent nucleoli has been described as resulting from nucleolar fusion, implying that nucleoli on individual p-arms have a propensity to coalesce (Anastassova-Kristeva 1977; Mangan et al. 2017). Critically however, we cannot distinguish between cause and effect. It is equally plausible that acrocentric p-arms have a propensity to self-associate, with formation of multivalent nucleoli being a consequence. Here, we use CRISPR/Cas9 mediated mega-base scale precision chromosome engineering (PCE) (Essletzbichler et al. 2014) to determine the chromosomal requirements for the participation of acrocentric p-arms in multivalent nucleoli. We reveal that rDNA arrays are not a prerequisite for their association with multivalent nucleoli, nor is there a requirement for the DJ or PJ. Instead, sequences further distal and proximal to NORs act redundantly to facilitate acrocentric gathering. Additionally, we demonstrate that there is no major contribution from the acrocentric centromere.

## Results and Discussion

### Sequences across acrocentric p-arms are nucleolar associated

Utilising a combination of rDNA and DJ probes, we had determined that in hTERT-RPE1 cells all 10 acrocentics p-arms are nucleolar associated, despite one of the chr22 haplotypes having little or no rDNA (van Sluis et al. 2020). To reveal the extent of acrocentric p-arm nucleolar association, we have designed a series of FISH probes that span the p-arms. An NOR probe includes the rDNA array as well BACs encompassing both the DJ and PJ. Combined hSat1 and βSat probes recognise distal regions preferentially. Finally, hSat3 was used to identify predominantly proximal regions. The distribution of sequences recognised by these regional probes on p-arms of the T2T CHM13 genome (Nurk et al. 2022) is shown in Fig. 1B. 3D-immunoFISH using these p-arm regional probes and an antibody to GC protein Nop52 (van Sluis et al. 2016) to visualise nucleoli was performed on both normal and Actinomycin D (AMD) treated RPE1 cells (Fig. 1B). At low concentrations AMD is a selective inhibitor of RNA polymerase I, inducing nucleolar segregation and more clearly delimiting nucleoli and PNH (Mangan and McStay 2021). This revealed that all 10 acrocentric p-arms are intimately associated along their length with nucleoli, consistent with the PNH being largely composed of acrocentric p-arms. Next, we performed PCE to elucidate the rules of engagement for acrocentric p-arms in multivalent nucleoli.

### Deletion of rDNA arrays from human acrocentrics in mouse A9 hybrids

To formally test the requirement for rDNA arrays, we sought to assess the nucleolar association potential of p-arms in which they had been deleted using CRISPR/Cas9. As DJ and PJ sequences are shared among the acrocentrics, we initially chose to perform the rDNA deletion on individual human chromosomes 14 and 15 (chr14 and chr15) held within mouse A9 mono-chromosomal hybrids A9-14 and A9-15 respectively (Cuthbert et al. 1995). The engineered chromosome was to be shuttled into human RPE1 cells by microcell mediated chromosome transfer (MMCT) as previously described (Meguro-Horike and Horike 2015; Mangan and McStay 2021). We designed a pair of gRNAs that would facilitate precise deletion of rDNA arrays from both chr14 and chr15 by CRISPR/Cas9. These gRNAs are located 804bp and ∼6.6kb distal and proximal to rDNA respectively (Fig. 2A). By co-transfecting A9-14 and A9-15 cells with pairs of Cas9-gRNA RNPs and screening for novel DJ/PJ fusions by PCR, we obtained two independent A9-14ΔhrDNA (#16 and #20) and two A9-15ΔhrDNA15 clones (#26 and #33) (Supplemental Fig. S1A). Sequencing revealed that DJ and PJ’s had fused with the deletion of a few additional bases, or in the case of one of the A9-15 clones (#33), inclusion of 65 bp of mouse sequence (Supplemental Fig. S1A). Deletion of rDNA was further confirmed by 3D-immunoFISH (Supplemental Fig. S1B).

**Figure 2.**
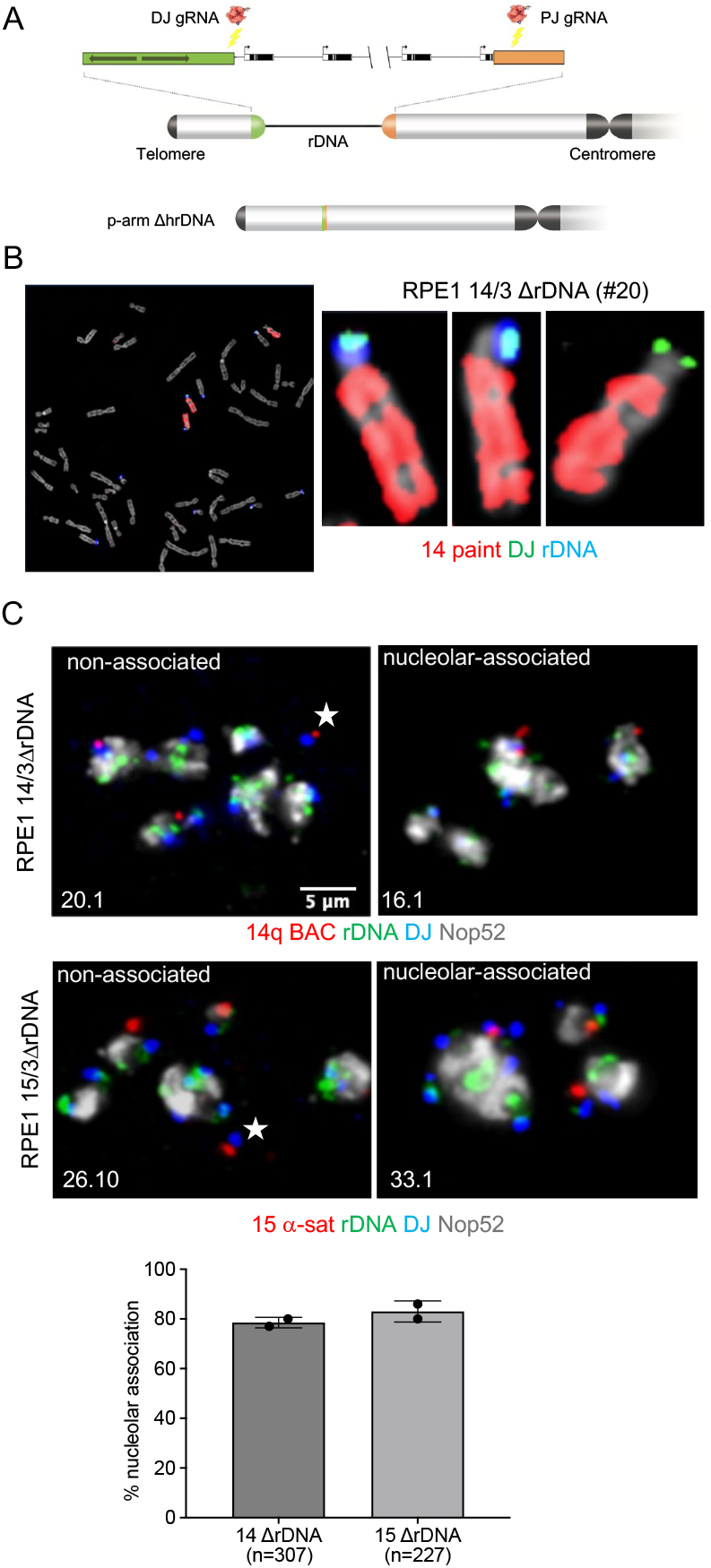
Deletion of rDNA from single human acrocentrics present in A9 mono-chromosomal hybrids. (A) Cartoon showing the CRISPR mediated rDNA deletion strategy and locations of DJ and PJ gRNAs. (B) M-FISH on RPE1 14/3 ΔrDNA with rDNA and DJ FISH probes and a chr14 paint. The three chr14s present in this cell line are shown on the right. Note the absence of rDNA on the transferred chr14 (14ΔrDNA). (C) 3D-immunoFISH on RPE1 14/3 ΔrDNA (upper panels) and RPE1 15/3 ΔrDNA (lower panels) containing 14ΔrDNA and 15ΔrDNA chromosomes from A9-14 and A9-15 respectively. Each panel represents a single nucleus. FISH probes included rDNA, DJ and either a chr14 q-arm BAC or α-sat15 and Nop52 antibodies. Representative images of cells in which the transferred chromosome was nucleolar associated or non-associated (indicated by an asterisk) are shown, with the clone identities indicated. Quantitation is shown below. Bars represent the mean ± standard deviation (SD) across both clones, *n* indicating the number of cells analysed.

### rDNA arrays are not required for association of acrocentric p-arms with multivalent human nucleoli

To test for the impact of rDNA deletion on nucleolar association potential in human cells, the engineered human acrocentrics were transferred into human RPE1 cells by MMCT as previously described (Mangan and McStay 2021). Successful chromosome transfer and the absence of carried over mouse chromosomes was confirmed by a combination of metaphase FISH (m-FISH) performed on spreads and STR (short tandem repeat) analysis (Fig. 2B and Supplemental Fig. S1C). We isolated two RPE1 clones containing an rDNA deleted chr14, one from each of the independent A9-14 ΔhrDNA clones (#16.1 and #20.1). Additionally, we isolated four RPE1 clones containing an rDNA deleted chr15, three from one A9-15 ΔhrDNA #26 (26.8, 26.9, 26.10) and one from #33 (33.1) (Supplemental Fig. S1C)

To assess the nucleolar association potential of the engineered chromosome, we performed 3D-immunoFISH on interphase cells, combining a chr15 specific α-satellite (α-sat15) centromere probe, DJ and human rDNA probes. ΔhrDNA15 could be distinguished from the endogenous chr15s by the absence of rDNA. Nucleoli were identified using antibodies to Nop52. To visualise ΔhrDNA14, we replaced a centromeric probe with a BAC that maps immediately proximal to centromere on the q-arm of chr14 (van Sluis et al. 2020). This was required as a chr14 centromere probe also recognises chr21, complicating the analysis. Combining the results from independent clones we determined that in 81.5% and 78.5% of cells the ΔhrDNA15 and ΔhrDNA14 respectively were observed to be nucleolar associated (Fig. 2C).

Next, we turned our attention to deleting endogenous rDNA arrays from acrocentrics present within standard RPE1 cells. While initially hesitant to do this, as the DJ and PJ gRNA sequences are present on all the acrocentrics, we were able to isolate a clone, RPE1-ΔrDNA13, in which the rDNA arrays present on both chr13 haplotypes had been deleted. rDNA deletion was confirmed by PCR which amplified the novel DJ/PJ junctions. Sequencing of 8 independent cloned PCR products revealed that both haplotypes have identical precise DJ/PJ fusions(Supplemental Fig. S2A). Loss of rDNA was further confirmed by m-FISH (Fig.3A and Supplemental Fig. S2B). This also confirmed that the other acrocentrics retained their normal rDNA content.

Previous analysis had revealed that in RPE1 cells both chr13s have moderate sized rDNA arrays (van Sluis et al. 2020). Thus we expected an observable decrease in rDNA copy number. In line with expectation, digital PCR confirms that the rDNA copy number per cell drops from ∼340 repeats in RPE1 cells to ∼270 repeats in RPE1-MrDNA13 cells, a drop of 20% (Fig.3B).

To visualise chr13s in RPE1-ΔrDNA13 interphase cells we combined a 13q centromere proximal BAC probe (van Sluis et al. 2020) with Nop52 antibodies (Fig. 3C). In 75% of RPE1-ΔrDNA13 cells we observe that one or both chr13s are nucleolar associated. This compares to 95.7% in standard RPE1 cells. We conclude that while rDNA arrays are not essential for participation of acrocentric p-arms in multivalent nucleoli they likely contribute. This is further supported by our previous studies with neoNORs, synthetic rDNA arrays on metacentric chromosomes, that can associate with endogenous multivalent nucleoli (Grob et al. 2014).

**Figure 3.**
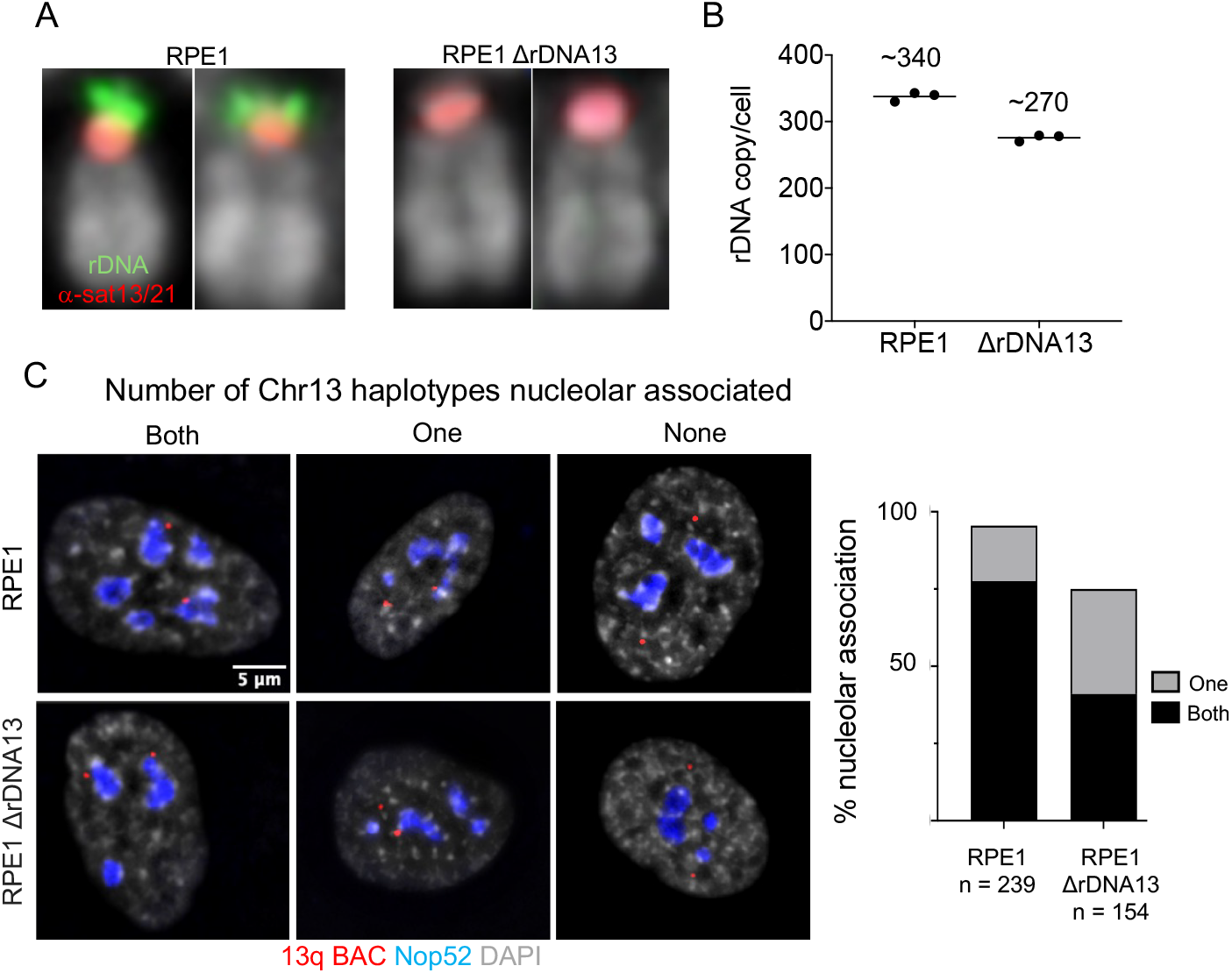
Deletion of endogenous rDNA arrays in RPE1 cells. (A) M-FISH was performed on normal RPE1 cells and RPE1-ΔrDNA13 with α-sat13/21, α-sat14/22 and rDNA FISH probes. Isolated chr13 pairs from each are shown. The full spreads are shown in Supplemental Fig. S2B. (B) rDNA repeat copy numbers in RPE1 and RPE1-rDNA13 cells were determined by digital PCR, performed in triplicate. The average repeat number for each cell type is shown above. (C) 3D-immunoFISH on RPE1 and RPE1-ΔrDNA13 with a 13q BAC FISH probe, and Nop52 antibodies. Representative images for cells in which both, one or no chr13s are associated with nucleoli are shown. Quantitation is shown on the right, with *n* indicating the number of cells analysed.

### Role of NOR distal and proximal sequences

Then we turned our attention to the role of NOR distal and proximal p-arm regions in formation of multivalent nucleoli. Initially we focused on the NOR distal region of chr15. We combined the distal gRNA used for rDNA deletion with a gRNA positioned approximately 32kb from the telomere of chr 15 (Fig. 4A). Notably, this telomeric guide appears unique to chr15 and is present within the T2T CHM13 genome (Nurk et al. 2022) as well as our A9-15 p-arm sequences (data not shown). This gRNA pair was used to generate RPE1-15Δdistal, in which the NOR distal region of one chr15 had been deleted, resulting in the telomere now being positioned ∼32kb from the rDNA array. PCR and DNA sequencing confirmed the presence of this novel junction (Supplemental Fig.3A). The two chr15 haplotypes of RPE1 cells can be distinguished based on their rDNA content and the extent of centromeric α-sat DNA (Supplemental Fig.4). One haplotype, labelled here as Hap1, has a relatively small rDNA array and a large α-sat array, while the other (Hap2) has a large rDNA array and a low amount of α-sat. The NORs on both are active as determined by the presence of UBF on metaphase chromosomes (van Sluis et al. 2020). Immuno-FISH performed on metaphase spreads from RPE1-15Δdistal revealed that the deletion had occurred on the high rDNA/low α-sat Hap2 and revealed the presence of UBF on the deleted chromosome Fig. 4B). Thus we can conclude that the rDNA array on the deleted chromosome is active, despite the absence of a DJ and the positioning of a telomere immediately distal.

**Figure 4.**
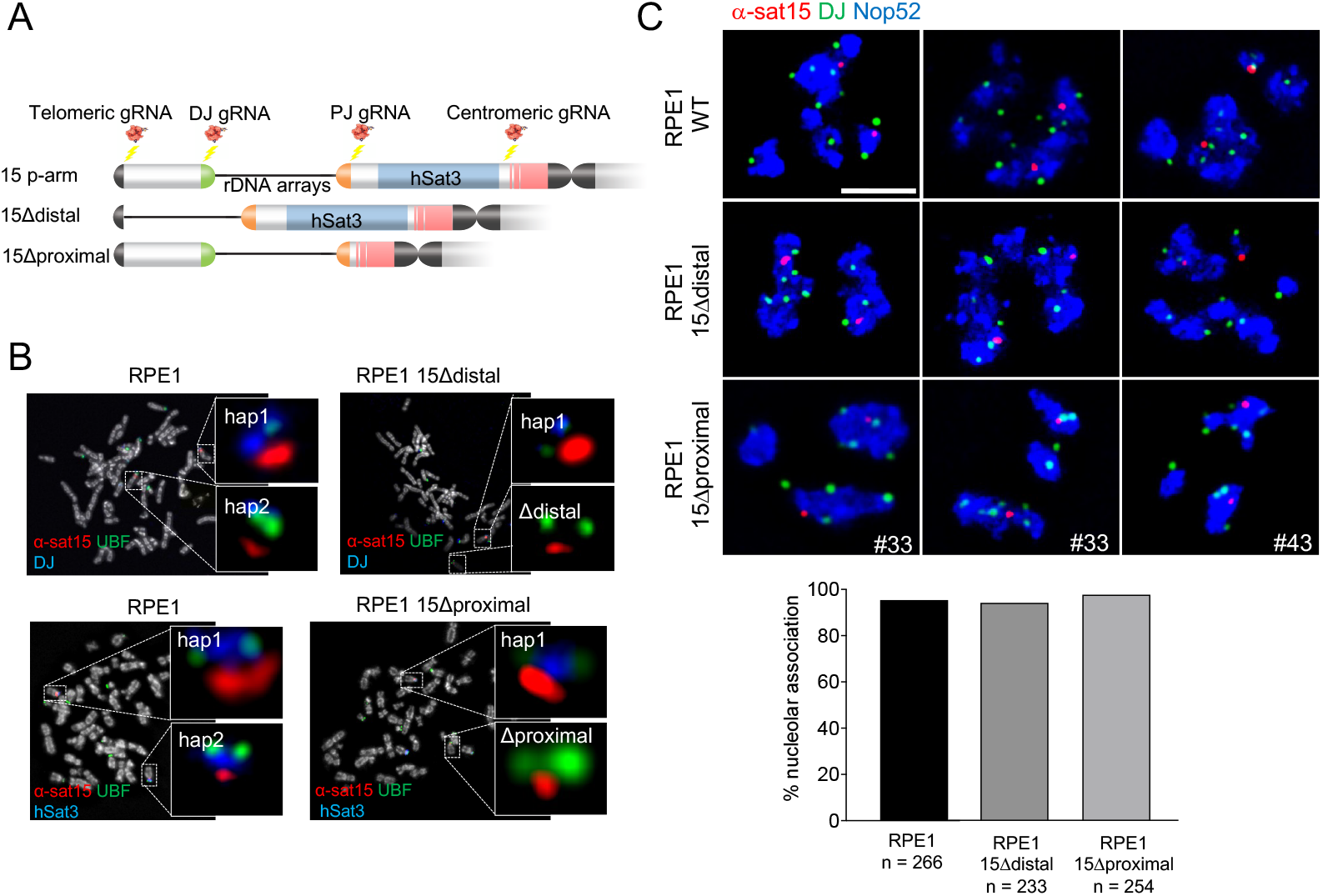
Consequences of deleting NOR distal or NOR proximal regions. (A) Cartoon showing the CRISPR mediated distal and proximal deletion strategies and location of relevant gRNAs. (B) Upper panels show immunoFISH of metaphase spreads from RPE1 and RPE1-15Δdistal cells probed with α-sat15 and DJ FISH probes and hUBF antibodies. Lower panels show immunoFISH on metaphase spreads from RPE1 and RPE1-15 proximal cells probed with α-sat15 and hSat3 FISH probes and hUBF antibodies. In each case the p-arm of both chr15 haplotypes are shown enlarged. (C) 3D-immunoFISH on RPE1 (upper panels), RPE1-15Δdistal (middle panels) and RPE1-15 proximal (bottom panels) cells with DJ and α-sat15 FISH probes and Nop52 antibodies. A single nucleus is shown in each panel, the clone names for both RPE1-15Δproximal clones are indicated and the scale bar represents 5mm. Quantitation of nucleolar association is shown below with *n* indicating the number of cells analysed. Note that for RPE1-15Δproximal cells, we have pooled data from clones #33 and #43.

To determine if this distal deletion impacted on the involvement of this chromosome in multivalent nucleoli, RPE1 and RPE1-15Δdistal interphase cells were probed with α-sat15, rDNA and DJ FISH probes and Nop52 antibodies. In standard RPE1 cells we observed that both chr15s are associated with a multivalent nucleolus, determined by the presence of multiple DJ signals, in 95% of cells. In RPE1-15Δdistal we observe that both 15s are associated with a multivalent nucleolus in 93% of cells. We can now conclude that the telomere to rDNA chromosome interval, including the DJ, is not essential for the participation of an acrocentric in multivalent nucleoli.

To assess the role of NOR proximal region of p-arms in formation of multivalent nucleoli, we sought to delete the rDNA to centromere interval from chr15 within RPE1 cells. We designed a chr15 specific gRNA that is positioned on the p-arm ∼1.7 Mb from the centromere in T2T CHM13 (Fig. 4A). This gRNA was similarly positioned on 15p in A9-15. The chr15 centromere gRNA was combined with the PJ gRNA used above to delete rDNA. This allowed us to generate two independent RPE1-15Δproximal clones (#33 and #43). Probing metaphase spreads from RPE1-15Δproximal clones revealed that in both, the high rDNA/low α-sat15 Hap2 had been targeted (Fig. 4B). Sanger sequencing of PCR products from independent clones confirmed they contained the novel junction (Supplemental Fig.3B). The deleted chromosomes were further shown to retain DJ signal and lack the major hSat3 array normally located proximal to rDNA. Finally, we could confirm that in 95% of spreads the rDNA array remained UBF loaded, thus active, despite proximity to the centromere.

3D-immunoFISH reveals that in WT RPE1 and RPE1 15Δproximal cells, both 15 p-arms are found associated with a multivalent nucleolus (multiple DJ signals) in 95% and 98% of cells respectively. Thus we conclude that the rDNA to centromere interval of chr15 is not essential for the participation of an acrocentric in multivalent nucleoli.

We have now demonstrated that neither NOR distal or proximal regions are individually essential for inclusion of acrocentric p-arms in multivalent nucleoli. Combining these findings with the lack of an rDNA requirement, demonstrated above, points to two contrasting hypotheses. The first posits that acrocentric p-arms have redundant features, both NOR distal and proximal, capable of promoting their association. An alternative hypothesis is that the signal for association lies elsewhere on the acrocentric chromosome. Published observations that centromeres associate with nucleoli, particularly in human ESCs, point to the possibility that ‘specialised’ acrocentric centromeres may be involved (Rodrigues et al. 2023; Yeo et al. 2026). In order to distinguish between these competing models, we next sought to delete an entire acrocentric p-arm, while leaving its centromere intact.

### Acrocentric p-arm deletion

Deletion of an entire acrocentric p-arm in RPE1 cells was achieved by combining the chr15 specific sub-telomeric and centromeric gRNAs utilised above (Fig. 5A). We isolated a clone, RPE1-15ΔP, in which the p-arm of the low rDNA/high α-sat15 Hap1 had been deleted. DNA sequencing showed the presence of unique telomere/centromere fusion (Supplemental Fig.5A). Further confirmation came from m-FISH which reveals that hSat3, rDNA, DJ and PJ had been deleted in Hap1ΔP (Fig. 5B and Supplemental Fig. 5B). Additionally, a telomere FISH signal now abuts the centromere on Hap1ΔP (Fig. 5C and Supplemental Fig.5C). We have not observed any chromosomal instability in this clone, indicating that juxtaposition of the telomere and centromere has no measurable impact on either of their functions.

**Figure 5.**
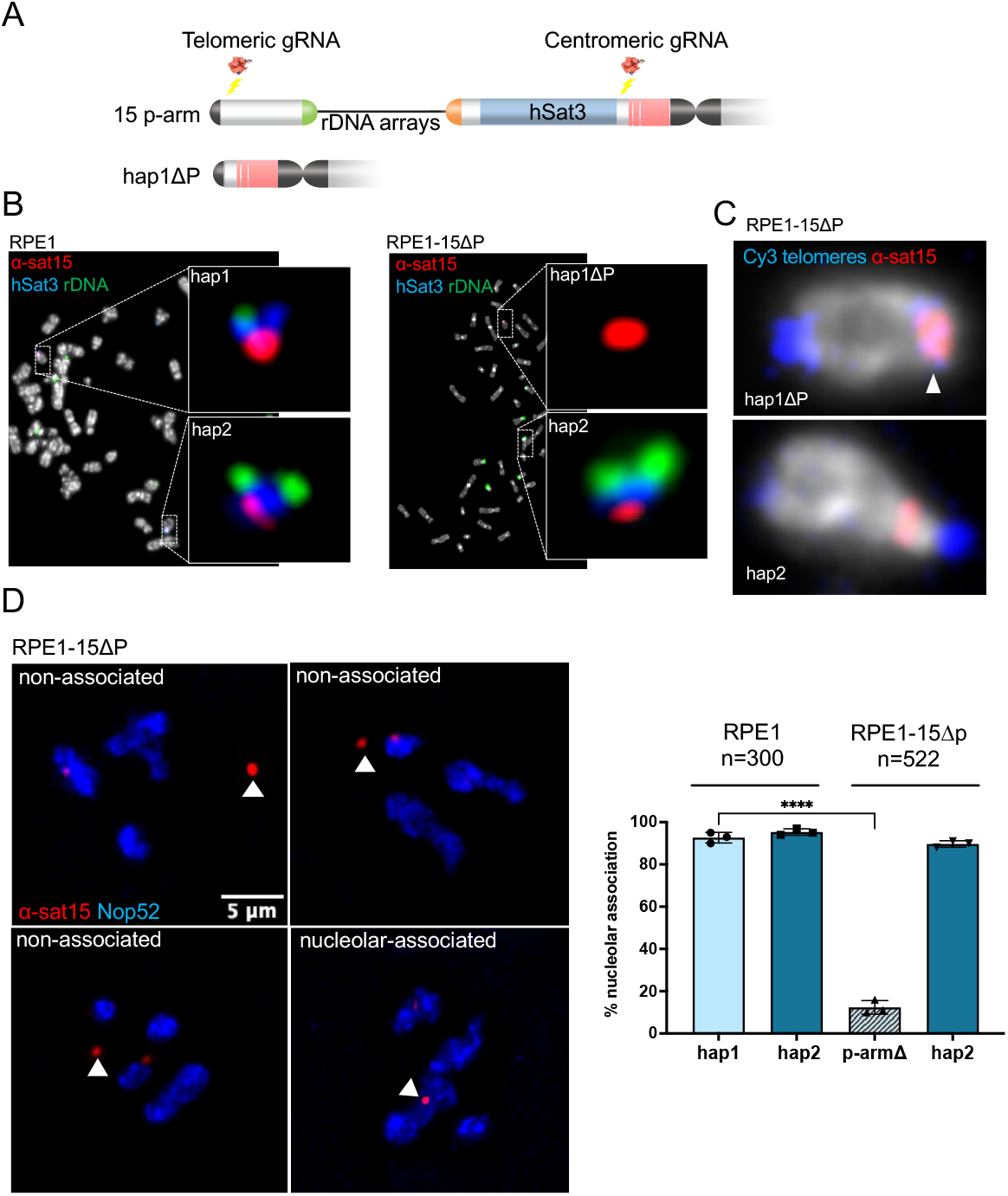
Deletion of full acrocentric p-arm. (A) Cartoon showing the CRISPR mediated p-arm deletion strategy and location of relevant gRNAs. (B) M-FISH on RPE1 and RPE1-15 P with α-sat15, hSat3 and rDNA probes. Both chr15 haplotypes for each cell type are enlarged. (C) FISH on RPE1-15 P metaphase spreads with a telomere (Cy3 labelled) and α-sat15 probes. Both chr15 haplotypes are shown. The arrow head indicates juxtaposed telomere and centromere signals on hap1 P. The full metaphase spread is shown in Supplemental Fig.5C. (D) 3D-immunoFISH on RPE1-15ΔP cells with α-sat15 and Nop52 antibodies. Each panel depicts a single nucleus and the position of the Hap1ΔP in each is indicated with an arrowhead. Quantitation of the nucleolar association for each chr15 haplotype is shown on the right, *n* indicating the number of cells analysed.

To address the impact of p-arm deletion on the chromosome positioning, normal RPE1 and RPE1-15ΔP cells were probed using α-sat15 and a Nop52 antibody. In normal RPE1 cells α-sat15 signals from Hap1 and Hap2 contact nucleoli in 95% and 98% of cells respectively (Fig. 5D). In contrast, within RPE1-15ΔP cells the Hap1ΔP centromere is dramatically repositioned, abutting nucleoli in only 12% of cells. The positioning of chr15 Hap2 remains unaltered. Thus we conclude, chromosomal features that facilitate gathering of acrocentric chromosomes leading to the formation of multivalent nucleoli are restricted to acrocentric p-arms, and do not involve either specialised acrocentric centromeres or q-arms.

Combined, the results from our PCE experiments provide compelling evidence that formation of multivalent nucleoli is primarily driven by the propensity of acrocentric p-arms to self-associate. Moreover, the chromosomal domains underlying this are redundant and distributed along the length of acrocentric p-arms.

Other than rDNA, the DJ and PJ are the only sequences that are shared in sequence and relative position among all five acrocentrics, yet neither of these are essential for acrocentric association. The distal deletion we generated includes most of the DJ. As systemic depletion of DJ encoded lncRNAs negatively impacts ribosome biogenesis (van Sluis et al. 2019; Ataei et al. 2025), we were surprised that rDNA, now abutting the telomere on the deleted chromosome, remained active. A possible solution to this paradox is that DJ transcripts, from intact acrocentrics, may function in trans, albeit within multivalent nucleoli, to regulate NORs.

NOR distal and proximal regions of acrocentric p-arms are composed largely of a number of satellite DNA classes and segmental duplications in various combinations and lengths (Nurk et al. 2022; McStay 2023; Guarracino et al. 2023). Thus, apart from rDNA, PJ and DJ, acrocentric p-arms are more thematically similar rather than displaying high sequence similarity along their lengths, heterochromatin being the obvious theme. While the primary aim of this work was identifying the underlying DNA that drives acrocentric associations, our findings point to a role for heterochromatin. The proliferation marker Ki-67, that spatially organises heterochromatin in early G1, possibly through its interaction with HP1, prior to it being sequestered within multivalent nucleoli, may be involved (Booth et al. 2014; Sobecki et al. 2016; Bizhanova and Kaufman 2021; van Schaik et al. 2022; Schichler et al. 2026).

While a number of vertebrates, for example Xenopus laevis and rat kangaroo (Protorous tridactylis) have a single NOR per chromosome set, most mammals have multiple NORs. Not only does this provide redundancy, as evidenced by the lack of phenotype observed in individuals carrying a Robertsonian translocation (Guarracino et al. 2023; de Lima et al. 2025), but it also provides greater opportunity for fixing chromosomal position within the nucleus. Thus in humans, the self-association of acrocentric p-arms to form multivalent nucleoli provides a major positional cue for at least the ten acrocentric chromosomes. 3D-immunoFISH with a DJ probe, acrocentric paints and Nop52 antibodies, demonstrates that acrocentric q-arms make minimal contact with the nuclear lamina (Supplemental Fig.6). As acrocentric associations are largely independent of rDNA, the correct positioning of acrocentrics with low or no rDNA, is ensured. This is nicely illustrated in the primary cell line CCD-1079Sk, fibroblasts that were isolated from the skin of a normal male neonate donor. Within these cells, four of the ten acrocentrics lack detectable rDNA yet are associated with multivalent nucleoli (van Sluis et al. 2020). Furthermore, these cells can be efficiently converted into iPS cells that maintain the developmental potential to differentiate into all three primary germ layers (Yu et al. 2007). There have also been reports that in certain cell lineages, chr1 and chr9 can associate with multivalent nucleoli (Stahl et al. 1976). Given our results, this is likely due to their extensive pericentric satellite DNA/heterochromatin. Finally, nucleolar association of non-acrocentric chromosomes can be more dynamic, such as when the inactive X chromosome in female cells is undergoing DNA replication (Zhang et al. 2007).

Previously we have argued that acrocentric p-arms are dedicated to ribosome biogenesis, with the chromosomal context of rDNA arrays ensuring both their genetic isolation and genomic stability (McStay 2023). Additionally, the p-arm derived PNH may provide physical separation of nucleoli from the remaining nucleoplasm. To these functions we suggest including positional cues for acrocentric chromosomes that are independent of their nucleolar roles.

## Materials and Methods

### Cell culture

A9 hybrids and hTERT-RPE1 cells were maintained as previously described (van Sluis et al., 2019). To induce nucleolar segregation, cells were treated with 0.1 µg/mL Actinomycin D (AMD) (Sigma) for 1 h. MMCT was carried out as previously described (Meguro-Horike and Horike 2015; Mangan and McStay 2021).

### Targeted p-arm chromosomal deletions

A near full T2T genome for RPE1 cells has recently been determined (Volpe et al. 2025). However, as the rDNA arrays and much of the acrocentric p-arms are missing, we utilised T2T CHM13 (Nurk et al. 2022) and our own data from A9 hybrids to design gRNAs (van Sluis et al. 2019). Production of gRNAs and CRISPR genome editing were performed as previously described (Mangan and McStay 2021). Briefly, 4 µg of each gRNA were precomplexed with 8.2 µg of Alt-R S.p. Cas9 nuclease V3 (Integrated DNA Technologies) for 10 min at room temperature. RNPs were combined and co-electroporated with 500 ng Puromycin mRNA into 100,000 cells using 1400V 10ms x 3 pulses for A9 cells, and 1350 V 20 ms x 2 pulses for RPE1 cells using a Neon transfection system (Thermo Fisher Scientific). 2 hours post-transfection, cells were selected with puromycin (Sigma) for 48 hrs, 2 µg/mL for RPE1 cells or 10 µg/mL for A9 cells. Cells were seeded for single-cell colonies. Following 14 days, single-cell colonies were picked into 24-well plates. 7 days later, individual colonies were screened using Phire Direct PCR Master Mix (Thermo Fisher Scientific) for the presence of novel junctions. Oligonucletide sequences used in gRNA production and PCR are presented in Supplemental Table S1.

### Fluorescent in situ hybridization (FISH) on cells and chromosome spreads

For description of all FISH probes see Supplemental Materials and Methods. 3D-immunoFISH and m-FISH were performed as previously described (Mais et al. 2005) (van Sluis et al. 2016). For m-FISH, fixed cells were dropped onto pre-chilled glass slides. For immunofluorescence on metaphase spreads, non-aged slides were fixed in 1% paraformaldehyde (PFA) (w/v) in 1x PBS for 10 min, and then subjected to FISH. UBF was immunostained post-hybridization, followed by three 15-min PBS washes. Slides were incubated with secondary antibodies (Jackson Immuno Research) for 1 h at 37 °C, followed by three 15-min PBS washes. Slides were mounted in VectorShield® (Vector Laboratories) with or without DAPI. All primary and secondary antibodies are listed in Supplemental Table S2. Telomere FISH is described in Supplemental Materials and Methods.

### Microscopy

All Images were captured using a Leica DMi8 microscope as previously described (van Sluis et al. 2019). To assess nucleolar association, α-sat15 was used for chr15. For chr13 and chr14, BAC probes were used (Supplemental Table S1). Distances to nucleolar boundary were measured from maximum intensity projections. Acrocentric chromosomes were scored as nucleolar-associated if α-sat/BAC signal overlapped with, or was positioned within 0.5 µm of nucleolar boundary. For RPE1 cells, nucleolar boundary was identified using an antibody to the granular component protein Nop52. For A9 hybrids, nucleolar boundary was identified either with Treacle or Nucleolin, as the human Nop52 antibody does not recognise mouse Nop52.

## Acknowledgments

We thank Thomas Durkin for performing digital PCR and Carol Duffy for manuscript editing. This work was supported by Wellcome Trust Investigator Award 223049/Z/21/Z to B.M and a PhD Scholarship from the Irish Research Council GOIPG /2021/852 to K.G.

## Author contributions

B.M. designed the research and wrote the manuscript. K.G. and H.M. performed the experiments and commented on the manuscript.

